# Unmyelinated neurons use Neuregulin signals to promote myelination of neighboring neurons in the CNS

**DOI:** 10.1101/2022.03.23.485365

**Authors:** Daniel E. Lysko, William S. Talbot

**Affiliations:** Department of Developmental Biology, Stanford University, Stanford, CA 94305

## Abstract

The signaling mechanisms neurons use to modulate myelination of circuits in the central nervous system (CNS) are only partly understood. Through analysis of isoform-specific *neuregulin1* (*nrg1*) mutants, we identify *nrg1* type II as an important regulator of myelination in the zebrafish CNS, required for normal myelination of two classes of spinal cord neurons. Surprisingly, *nrg1* type II reporter expression is prominent in unmyelinated Rohon-Beard (RB) sensory neurons, while myelination of interneurons controlling the escape response circuit is reduced in *nrg1* type II mutants. Cell type-specific loss-of-function studies indicate that *nrg1* type II is required in RB neurons to signal to other neurons, not oligodendrocytes, to modulate spinal cord myelination. Together, our data support a model in which unmyelinated neurons express Nrg1 type II proteins to regulate myelination of circuit partners, a mode of action that may coordinate function of circuits in the CNS involving both unmyelinated and myelinated neurons.

**Summary points:**

1. *nrg1* type II is required for normal myelination of diverse neuronal classes in the zebrafish spinal cord
2. Surprisingly, *nrg1* type II reporter expression is prominent in unmyelinated Rohon-Beard neurons
3. Cell type-specific knockdown indicates that myelination of CoPA neurons requires *nrg1* type II function in unmyelinated Rohon-Beard neurons
4. The Nrg1 receptor *erbb2* is required in neurons, but not oligodendrocytes, for normal myelination

## Introduction

The staggering complexity of neuronal circuits in the central nervous system requires precise regulation of axonal conduction to coordinate the transmission of action potentials. Emerging evidence indicates that modulation of myelination can tune axonal conduction velocity to coordinate circuit activity, thereby enabling coherent input, output, and behavior (Almeida and Lyons, 2017; Chang et al., 2016; Etxeberria et al., 2016; Stanford, 1987; Waxman, 1997). Specific patterns of myelin sheaths have been described on distinct neurons in the brain and spinal cord (Call and Bergles, 2021; Koudelka et al., 2016; Tomassy et al., 2014; Zonouzi et al., 2019), but the mechanisms that encode these patterns are only starting to be understood. Increasing evidence shows that myelination can be modulated by neuronal activity (Bacmeister et al., 2020; Gibson et al., 2014; Hines et al., 2015; Koudelka et al., 2016; Yang et al., 2020) as well as a wide variety of signaling molecules, including PDGF, BDNF, ATP, and Neuregulin, among others (Falls, 2003a; Hughes, 2021; Welsh and Kucenas, 2018), but our understanding of how these factors coordinate myelination within complex neuronal assemblies remains incomplete.

Neuregulin1 (Nrg1) signals have a well-defined role in peripheral nervous system (PNS) myelination (Birchmeier and Bennett, 2016; Brinkmann et al., 2008; Meyer and Birchmeier, 1995; Michailov et al., 2004; Nave and Salzer, 2006; Salzer, 2012), but their role in central nervous system (CNS) myelination is less clear. The human *NRG1* gene encodes more than 30 isoforms that act as extracellular ligands, binding to ErbB family receptor tyrosine kinases on the surface of target cells (Falls, 2003b; Mostaid et al., 2016; Steinthorsdottir et al., 2004). In the PNS, the transmembrane isoform Nrg1 type III is the key axonal signal that controls the proliferation, migration and myelination of ErbB-expressing Schwann cells (Lyons et al., 2005; Meyer and Birchmeier, 1995; Perlin et al., 2011). In CNS myelination, the function of the Nrg1 type III isoform is much more limited; myelin is present but reduced in the brain of ErbB receptor mutants (Makinodan et al., 2012; Taveggia et al., 2008). However, these and other studies (Brinkmann et al., 2008) report normal myelination in the spinal cord in Nrg1 and ErbB mutants, despite evidence from pioneering *in vitro* and *ex vivo* studies (Bermingham-McDonogh et al., 1997; Marchionni et al., 1993) that established Nrg1 signals as regulators of development, migration, or proliferation of oligodendrocytes, the myelinating cells of the CNS. The roles of other Nrg1 isoforms in CNS myelination are not well-known, although different Nrg1 isoforms regulate synapse formation, heart development, and other processes (Liu et al., 2010; Loeb et al., 1999; Nave and Salzer, 2006).

To investigate the roles of different Nrg1 isoforms in the CNS, we created a series of isoform-specific *nrg1* zebrafish mutant lines and analyzed myelination in the spinal cord using live imaging. Our results indicate that the Nrg1 type II isoform modulates the myelination of distinct neuronal classes in the spinal cord. Nrg1 type II reporter expression and cell type-specific CRISPR experiments provide evidence that unmyelinated sensory neurons expressing Nrg1 type II control the myelination of interneurons that mediate the escape response. Our results indicate that, in contrast to the PNS, Nrg1 type II signals through ErbB2 receptors on neurons, rather than oligodendrocytes, to modulate myelination in the CNS.

## Results

### *nrg1* type II is required for normal myelination in the spinal cord

The *nrg1* gene generates many isoforms through multiple promoters and alternative splicing. A unique promoter and 5’ exon distinguish each main class of isoforms, resulting in distinct tissue-specific expression patterns and N-terminal protein sequences: types I-VI are present in humans, while types I, II, and III are common to all vertebrates (Mostaid et al., 2016; Steinthorsdottir et al., 2004). To determine which Nrg1 isoforms have essential functions in myelin sheath formation in the CNS, we used CRISPR/Cas9 to create a series of mutations in the zebrafish *nrg1* gene. We used a gRNA targeting the common EGF domain to eliminate the function of all *nrg1* isoforms and gRNAs targeting each isoform’ s unique exons to create isoform-specific mutants (Figure 1A,B). To broadly assess myelination in these mutants we performed *in situ* hybridization on whole mount embryos for *myelin basic protein* (*mbp*), a marker of myelinating glia. In the PNS, animals homozygous for mutations in the EGF exon or type III-specific exon lacked *mbp* expression, in accord with previous studies of zebrafish and mouse mutants (Meyer and Birchmeier, 1995; Perlin et al., 2011), whereas *mpb* expression in the PNS appeared normal in the type I and type II homozygous mutants (Figure S1). All *nrg1* mutants expressed *mbp* in the CNS, consistent with previous analyses (Brinkmann et al., 2008; Perlin et al., 2011). To analyze myelin sheath formation at higher resolution, we visualized oligodendrocytes and their myelin sheaths using the transgenic reporter cldnk:GFP-CAAX in the *nrg1* EGF and isoform-specific mutants (Figure 1C). We analyzed myelin sheath formation in the dorsal spinal cord at 3.5 dpf, when individual oligodendrocytes can be imaged. There was a significant reduction in the total length of myelin sheaths produced by oligodendrocytes in both the EGF and type II-specific mutants, but there was no significant difference in either the type I-or type III-specific mutants (Figure 1D). There was no significant reduction in dorsal oligodendrocyte number in either the EGF or type II-specific mutants (Figure S1). In contrast to *nrg1* type III’ s requirement in the PNS, these results indicate that *nrg1* isoform type II is essential for normal myelination in the developing spinal cord.

**Figure 1.**
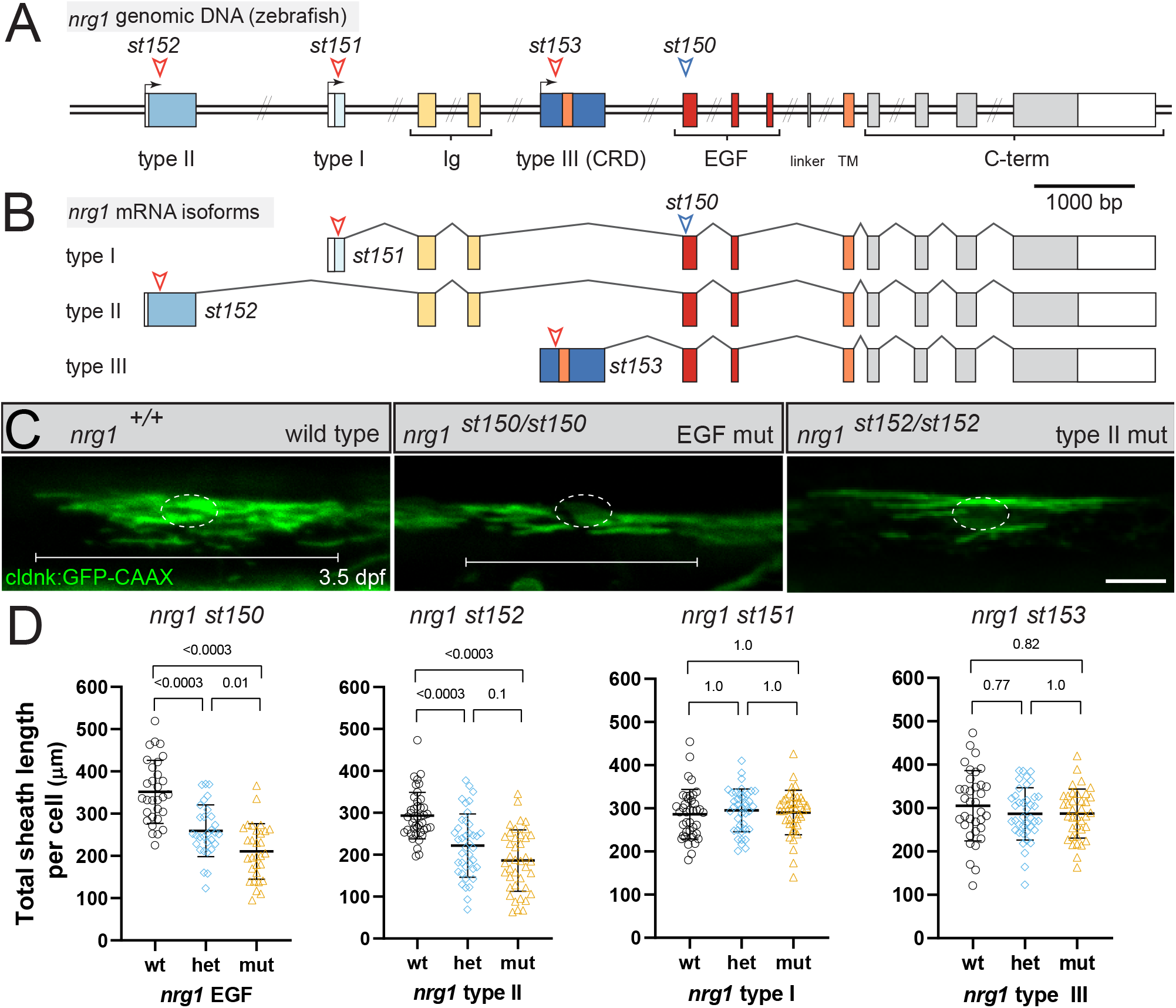
*nrg1* type II signals are required for normal myelination in the spinal cord. (A) Diagram of the zebrafish *nrg1* locus. Colored rectangles indicate coding exons, white rectangles indicate UTRs. Black arrows indicate alternative transcriptional start sites. Exon length is to scale, double slashes indicate intronic regions that are not to scale. Red arrowheads indicate gRNA targets in 5’ isoform-specific exons and allele location. Blue arrowhead indicates gRNA target in the EGF domain (st150 allele location) predicted to eliminate function of all *nrg1* isoforms. (B) Diagram of the mRNA splicing of *nrg1* isoforms, illustrating the isoform-specific nature of 5’ exons and chosen gRNAs. (C) Confocal images of oligodendrocytes in the dorsal spinal cord labeled using the claudink:GFP transgene. Wildtype fish have normal sheath parameters, while oligodendrocytes (OL) in EGF and type II mutants have fewer, shorter sheaths. Dotted circle indicates the OL cell body, while brackets indicate the breadth of a single OL. Scale bar, 20 μm. (D) Quantification of total myelin sheath length produced by individual oligodendrocytes in *nrg1* EGF and isoform-specific lines. OLs in *nrg1* EGF and type II mutants (mut) make significantly less myelin per OL than in wildtype (wt) animals, while no difference is observed in *nrg1* type I or III mutant animals. Error bars represent mean ± SD. A t-test was used to assess significance and p-values adjusted for multiple comparisons. Each point represents one OL from ≥ 4 animals per category; ≥ 30 OLs per category, one representative experiment shown from three replicates. Figure 1A,B modified from (Lysko et al., 2022).

### *nrg1* type II is required for normal myelination of diverse neuronal classes

In the larval zebrafish spinal cord, the majority of myelinated axons belong to three neuronal classes-- CoPA (commissural primary ascending), CiD (circumferential descending), and RS (reticulospinal) neurons (Koudelka et al., 2016). CoPA interneurons send ascending projections along the dorsal spinal cord to the hindbrain, while RS neurons project from the hindbrain to targets within the dorsal and ventral spinal cord (Higashijima et al., 2004; Lewis and Eisen, 2003). CiD interneurons send descending projections to targets within the dorsal spinal cord. To assess the myelination of these neuronal classes, we visualized the transgenic contactin fusion protein myelin reporter (Koudelka et al., 2016) in neurons in *nrg1* mutants. Transmembrane contactin (cntn) is localized to the axonal membrane, where it becomes excluded from ensheathed areas of the axon upon glial contact, revealing the “footprints” of myelin sheaths (Figure 2 A,B; green segments). We expressed the contactin reporter using pan-neuronal drivers in transient transgenic experiments to randomly label spinal cord neurons. Neurons expressing the reporter were classified by their characteristic positions, morphologies, and projections, allowing us to analyze the myelination of the major myelinated neuronal classes in *nrg1* mutants. In *nrg1* type II mutants, CoPA and reticulospinal neurons were myelinated significantly less frequently than in wildtype siblings (Figure 2A,B,C). Typical myelin sheath patterns were evident on many CoPA and reticulospinal neurons that were myelinated in *nrg1* type II mutants, indicating that other regulators of neuronal class-specific myelination remain intact in these mutants. These results indicate that *nrg1* type II regulates the myelination of both CoPA and reticulospinal neurons, the most highly myelinated classes of neuron in the spinal cord.

**Figure 2.**
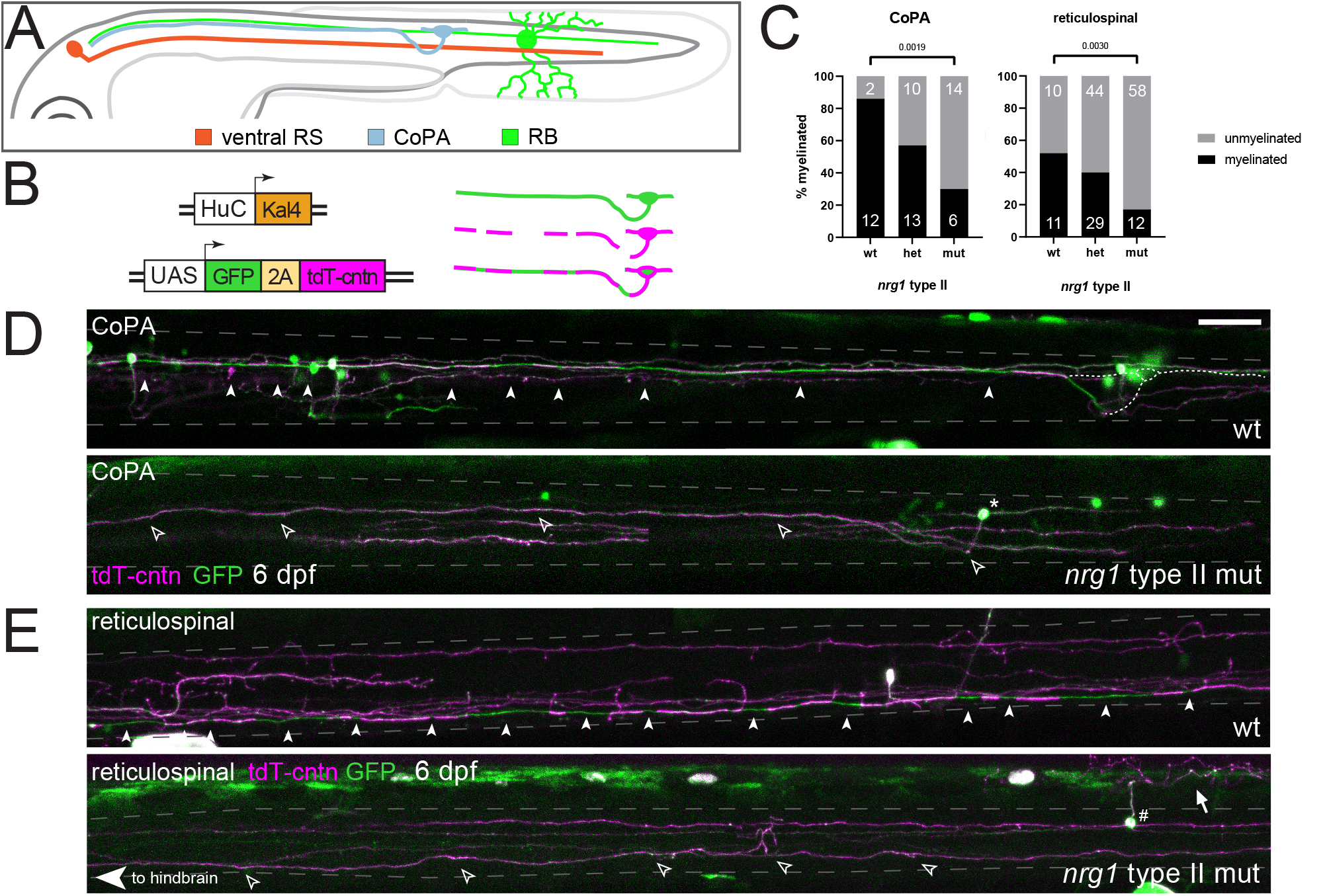
*nrg1* type II is required for normal myelination of diverse neuronal classes. (A) Illustration of spinal cord neurons assessed with the myelin reporter. RS neuron cell bodies reside in the hindbrain and project posteriorly toward the tail (orange). CoPA neurons reside in the dorsal spinal cord and project anteriorly to the hindbrain (blue). RB sensory neurons reside in the dorsal spinal cord, receive touch stimulation via their sensory arbor, and project both anteriorly to the hindbrain and posteriorly (green). (B) UAS:GFP-2A-tdTomato-contactin myelin reporter construct expresses GFP along the entire axon allowing neuronal type characterization, while tdTomato-contactin is excluded from ensheathed segments of the axon, allowing assessment of myelination status; thus myelin sheaths appear green in merged images. (C) Quantification of myelination status in CoPA and RS neurons. Percent of neurons that are myelinated indicated by bars, inset numbers indicate number of neurons assessed as myelinated or unmyelinated. Fisher’ s exact test was used to assess significance. (D) Myelin sheath pattern comparison between wild-type and *nrg1* type II mutant CoPA neurons. Arrowheads indicate myelin sheaths along the CoPA subject neuron. Dotted white lines indicate the position of CoPA cell body on the contralateral side of the spinal cord, which is not shown in this image projection. Asterisk indicates CoPA cell body in *nrg1* type II mutant image, while open arrowheads indicate the unmyelinated axon. Dashed grey lines indicates dorsal and ventral bounds of the spinal cord. (E) Myelin sheath pattern comparison between wild-type and *nrg1* type II mutant RS neurons. Arrowheads indicate myelin sheaths along the subject wildtype RS neuron, while open arrowheads indicate an unmyelinated RS axon in a *nrg1* type II mutant. A dorsal unmyelinated RB neuron (cell body, #; sensory arbor, arrow) sends projections posteriorly and also anteriorly to the hindbrain. Scale bar, 50 μm.

### Unmyelinated RB sensory neurons express *nrg1* type II reporter transgene

In the PNS, the level of Nrg1 type III expressed by a neuron regulates its myelination (Michailov et al., 2004; Taveggia et al., 2005). By analogy to the PNS, our analysis of myelination in the spinal cord of *nrg1* type II mutants raised the possibility that myelinated neurons might express Nrg1 type II signals, perhaps at different levels corresponding to their myelination rates. *nrg1* type II expression has been detected by *in situ* hybridization in spinal cord neurons (Honjo et al., 2008), but it was not possible from these studies to determine which neuronal classes were labelled. To examine *nrg1* type II expression with cellular resolution, we constructed a transgenic reporter with 3 kb of sequence upstream from the *nrg1* type II coding sequence driving expression of membrane-bound GFP (Figure 3A). Confocal microscopy of stable transgenic embryos revealed prominent expression of GFP in Rohon-Beard (RB) sensory neurons (Figure 3B-E), but no GFP expression in CoPA, CiD or RS neurons. RB neurons are specified early in neurogenesis (Figure 3B), and they have large somas located in the dorsal spinal cord. In the embryo, RB neurons form characteristic sensory arbors (arrow, Figure 3C), and they send descending projections towards the tail and ascending projections to the hindbrain (Figure 3E) via a dorsal axonal tract (Figure 3D) called the dorsal longitudinal fasciculus (Kaji and Artinger, 2004; Ogino and Hirata, 2018). Interestingly, RB neurons are typically unmyelinated (Koudelka et al., 2016). RB axons expressing the *nrg1* type II reporter pass near oligodendrocytes and axons in the dorsal spinal cord (Figure 3F), but course up to 50 micrometers away from oligodendrocytes and axons in the ventral spinal cord. In *nrg1* type II mutants labeled randomly with the contactin myelination reporter, we observed no significant change in the number of RB neurons (wt: 19 RBs; mut: 21 RBs per 100 fish analyzed) or their myelination status (wt, 96% unmyelinated; mut, 91% unmyelinated; p=0.64, Fisher’ s exact test). In addition, *nrg1* type II mutant larvae are responsive to trunk touch stimuli, a reaction dependent on the function of RB neurons (Drapeau et al., 2002; Ogino and Hirata, 2018). These data indicate that *nrg1* type II is not required for RB development or myelination, but is instead required for the normal myelination of both neighboring and distant axons that may not themselves express *nrg1* type II.

**Figure 3.**
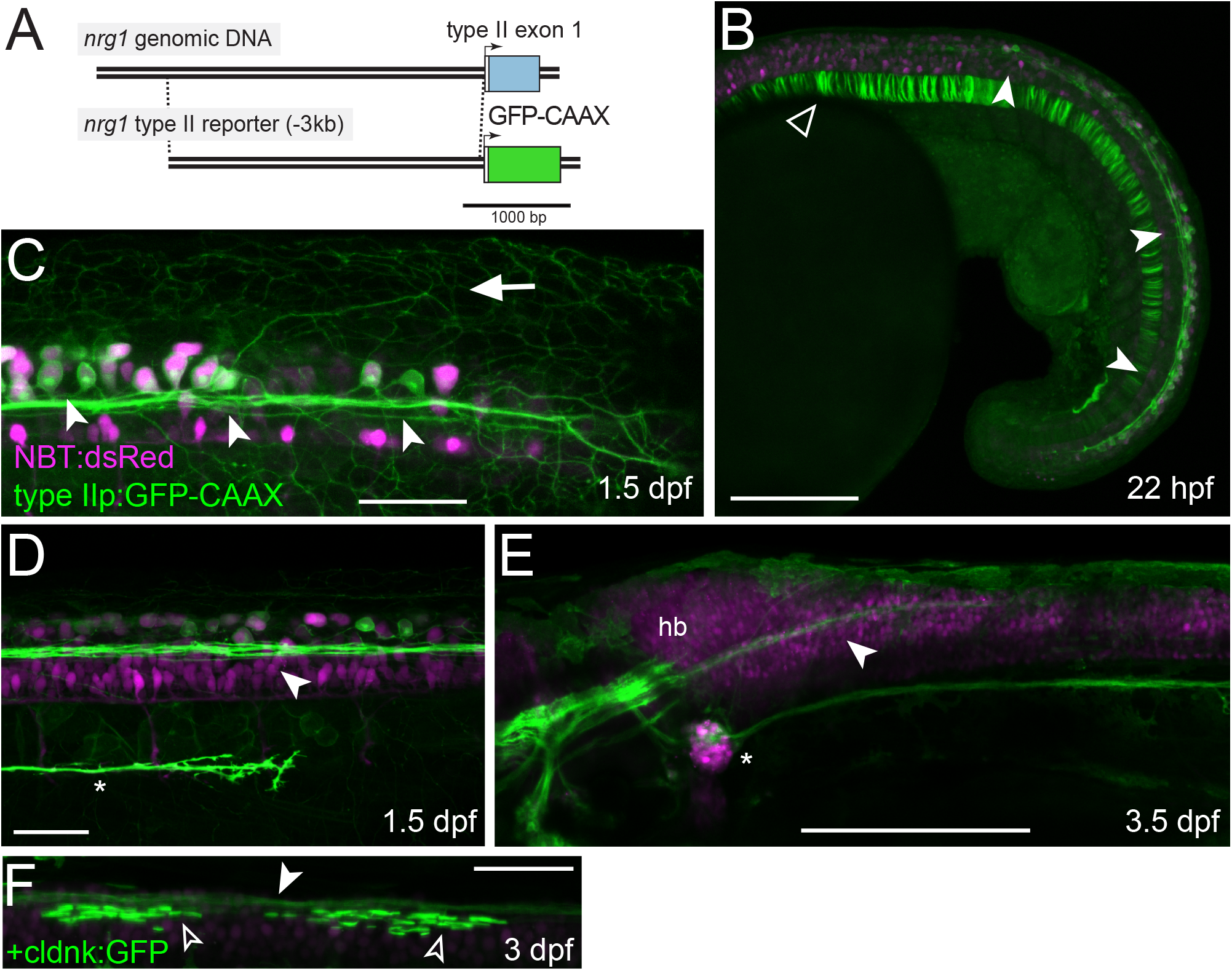
Unmyelinated RB sensory neurons express *nrg1* type II reporter transgene. (A) Diagram indicating location of 3kb regulatory element and reporter construct design. Colored rectangles indicate coding regions, white rectangles indicate UTRs. Arrows indicate transcriptional start sites. (B) *nrg1* type II reporter is expressed prominently in early-born Rohon-Beard (RB) neurons (arrowheads) and transiently in the notochord (open arrowhead) at 22 hours post fertilization. Scale bar, 200 μm. (C) *nrg1* type II reporter is expressed in neurons with large cell bodies and sensory arbors (arrow) characteristic of RB neurons at 1.5 days post fertilization (1.5 dpf). (D) Large, GFP+ RB neurons send axons (arrowhead) anterior and posteriorly via the dorsal longitudinal fasciculus (DLF). Reporter expression also occurs in the peripheral posterior lateral line nerve, which is growing posteriorly (asterisk). (E) GFP+ axons in the DLF (arrowhead) project anteriorly into the hindbrain (hb). GFP+ lateral line nerve descends from the posterior lateral line ganglion (*). Scale bar, 200 μm. (F) GFP+ axons from RB neurons (arrowhead) are closely apposed to oligodendrocytes in the dorsal spinal cord (open arrowheads). Scale bars, 50 μm unless indicated.

### *nrg1* type II is required in RB neurons to regulate myelination of other neurons in the spinal cord

To determine if *nrg1* type II is required in RB neurons to regulate myelination in the spinal cord, we adopted a cell-type specific CRISPR approach (Ablain et al., 2015; Chen et al., 2020; Marshall-Phelps et al., 2020), in which a gRNA is expressed ubiquitously and Cas9 is expressed in a cell type of interest. The *neurogenin1* gene regulates a transcriptional network necessary for RB specification (Andermann et al., 2002; Rossi et al., 2009), and a previous study identified a *neurogenin1* regulatory element termed LSEC that is mainly expressed in RB neurons (Blader et al., 2003), with a very similar spatiotemporal expression as our *nrg1* type II reporter. Consistent with the original study (Blader et al., 2003), we found that fish injected with an LSEC-GFP reporter transgene (Figure 4A) have prominent GFP expression in RB neurons (99% of GFP-expressing spinal cord neurons are RBs at 1 dpf; 94% at 2 dpf, 20 fish analyzed). We observed some LSEC-GFP expression in RS neurons (1% of neurons at 1 dpf, 6% at 2 dpf, rising to ∼50% by 3 dpf), rare expression in CoBL neurons and radial glia (<1%), and no expression in CiD or CoPA neurons, confirming the specificity of the LSEC regulatory element for RB neurons.

**Figure 4.**
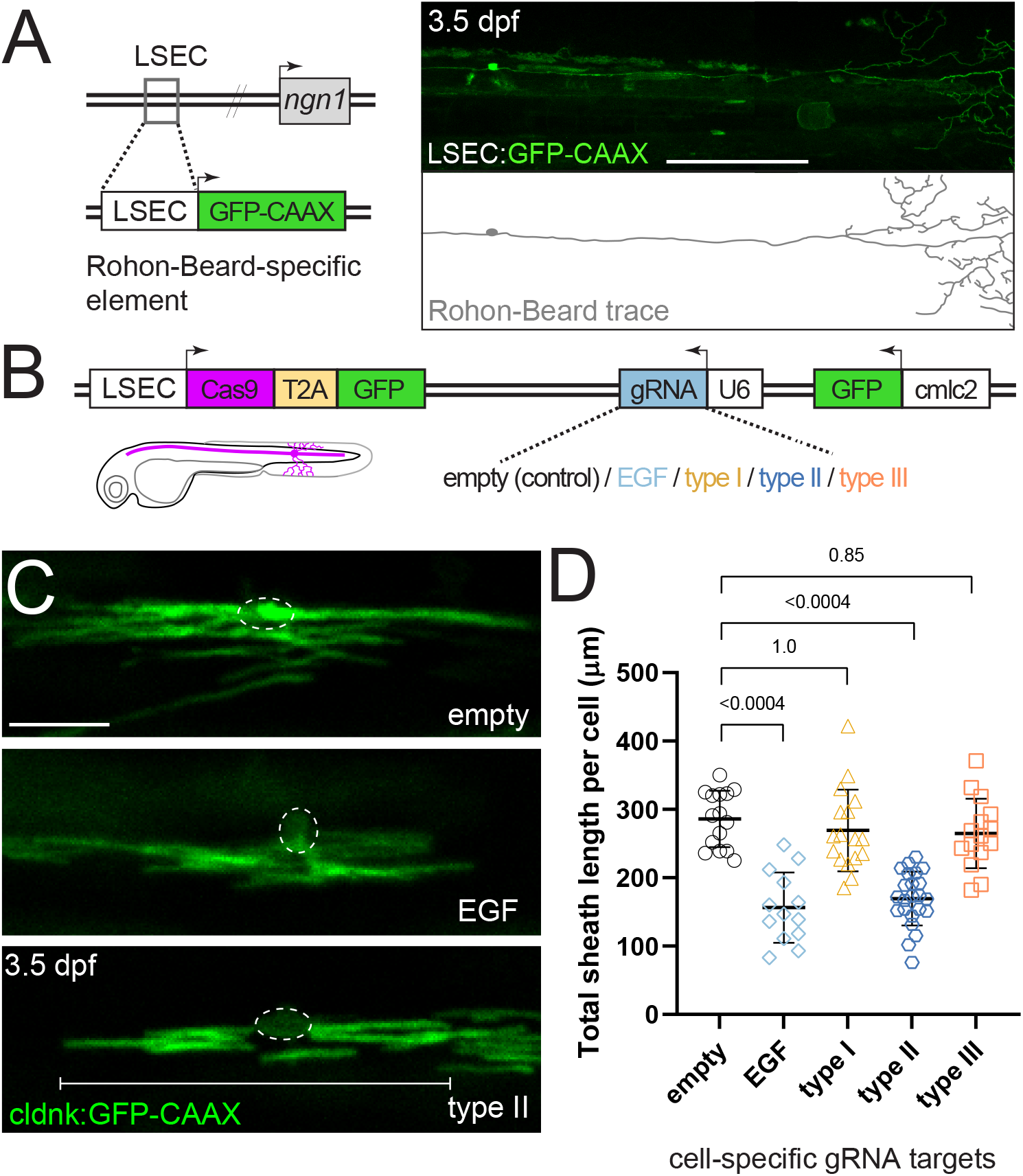
*nrg1* type II signals are required in RB neurons for normal myelination in the spinal cord. (A) Illustration showing the location of the LSEC upstream of the *neurogenin1* gene. In transient transgenic experiments, LSEC:GFP shows prominent expression in RB neurons with characteristic sensory arbors. Scale bar, 200 μm. (B) Diagram of cell type-specific CRISPR construct design, allowing RB expression of Cas9, and ubiquitous expression of EGF or isoform-specific *nrg1*-targeted gRNAs. (C) Confocal images of oligodendrocytes (OL) in the dorsal spinal cord labeled using the claudink:GFP transgene. Dotted circle indicates the OL cell body, while brackets indicate the breadth of a single OL. Scale bar, 20 μm. (D) Quantification of total myelin sheath length produced by individual OLs in animals expressing RB-specific Cas9, with indicated *nrg1* EGF and isoform-specific gRNAs. Error bars represent mean ± SD. A t test was used to assess significance and p-values adjusted for multiple comparisons. Each point represents one OL from ≥ 3 animals per category; one representative experiment shown from three replicates.

To determine if *nrg1* type II function is required in RB neurons for myelination of other neuronal classes, we used the LSEC element to express Cas9 in RB neurons, while simultaneously expressing efficient guide RNAs targeting the common or isoform-specific exons of *nrg1* (Figure 4B). We injected these constructs into wildtype fish expressing cldnk:GFP and measured dorsal oligodendrocyte myelin sheaths, as in our analysis of stable *nrg1* mutant lines. We observed normal amounts of myelin per oligodendrocyte in animals injected with a negative control construct and in animals injected with constructs targeting *nrg1* type I and III in RB neurons. In contrast, there was a significant reduction in myelin per oligodendrocyte in fish injected with constructs targeting *nrg1* type II-specific sequences and the common EGF exon in RB neurons (Figure 4C, D). These results demonstrate that *nrg1* type II function is required in RB neurons for myelination of other neuronal classes in the developing spinal cord.

### *erbb2* receptor is required in neurons, but not oligodendrocytes, for normal myelination in the spinal cord

Our data suggest that myelination of CoPA and RS neurons is regulated by Nrg1 type II expressed in RB neurons. To determine which cells may be responding to Nrg1 type II, we sought to disrupt ErbB receptor tyrosine kinases, which are activated by Nrg1 signals. We have previously characterized the PNS in zebrafish *erbb2* ^st61^mutants, in which Schwann cell migration and myelination is blocked (Lyons et al., 2005; Perlin et al., 2011). To determine if *erbb2* is essential for myelination in the CNS, we used transient transgenesis with an mbp:GFP-CAAX construct to label individual oligodendrocytes in the spinal cord of *erbb2*^st61^ mutants. Similar to the phenotype in *nrg1* EGF and type II mutants, there was a significant reduction in oligodendrocyte myelin sheath length in *erbb2* mutants at 5 dpf (Figure 5A). We used the cell-specific CRISPR approach to test if *erbb2* function was required in neurons or oligodendrocytes. We expressed an *erbb2* sgRNA, while simultaneously expressing Cas9 either pan-neuronally, in RB neurons with the LSEC element as in Figure 4, or in oligodendrocytes (Figure 5B). Interestingly, there was a significant reduction in myelin per oligodendrocyte in fish injected with the pan-neuronal construct targeting *erbb2*, but not with the constructs driving Cas9 expression in RB neurons or oligodendrocytes (Figure 5C, D). Together these data indicate that Nrg1 type II produced by RB neurons signals through ErbB2 in other neurons to regulate spinal cord myelination.

**Figure 5.**
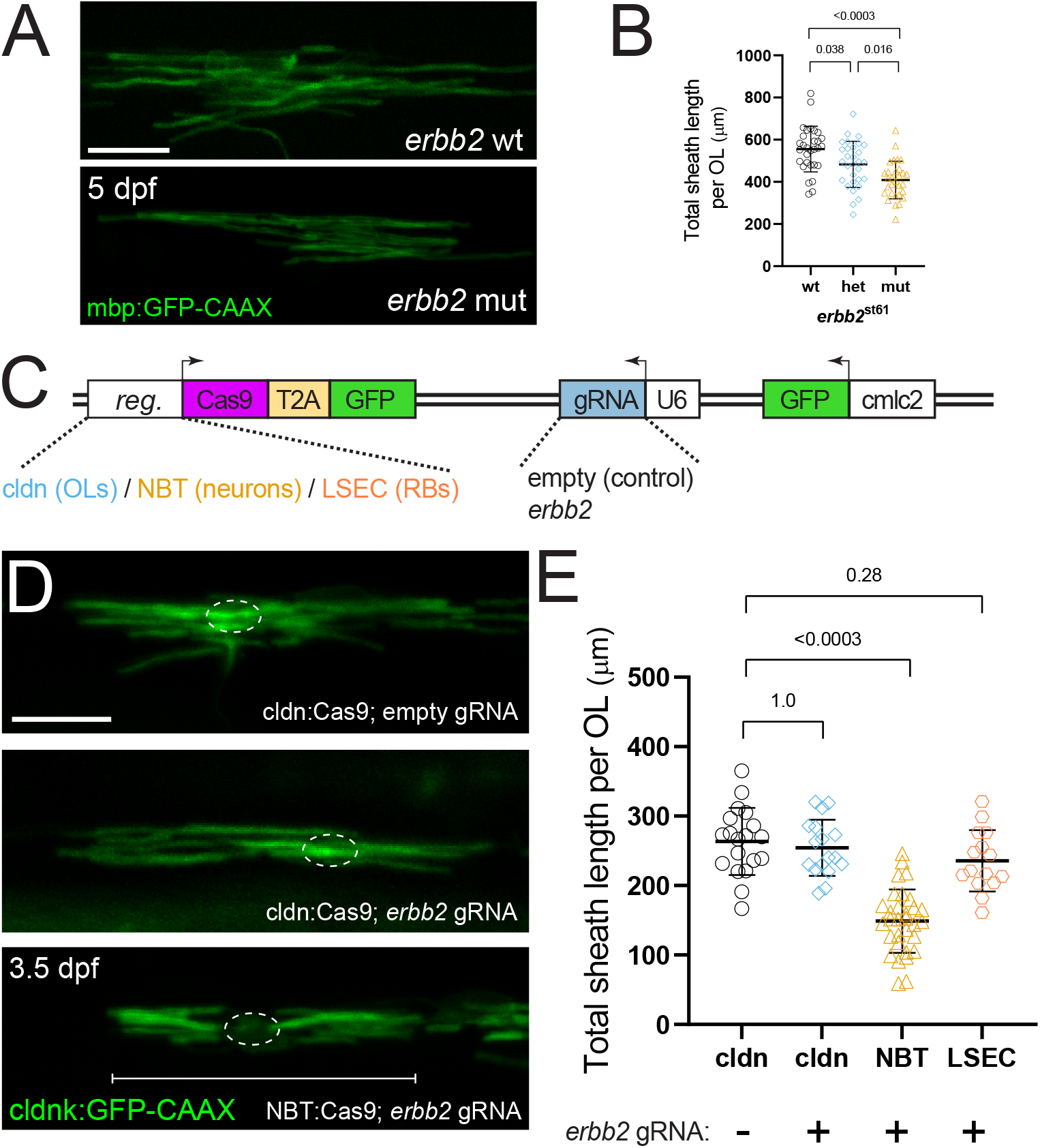
*erbb2* receptor is required in neurons, but not oligodendrocytes, for normal myelination in the spinal cord. (A) Confocal images of mbp:GFP-labeled oligodendrocytes (OL) in fish lacking *erbb2* generate less myelin than wildtype controls. Scale bars, 20 μm. (B) Quantification of myelin produced by individual OLs in animals wildtype, heterozygous, or mutant for *erbb2*^st61^. OLs in animals lacking *erbb2* produce significantly less myelin. (C) Diagram of cell type-specific CRISPR construct design, allowing expression of Cas9 in OLs (claudin), all neurons (NBT), or RBs (LSEC), and ubiquitous expression of *erbb2*-targeted gRNA. (D) Confocal images of oligodendrocytes in the dorsal spinal cord labeled using the claudink:GFP transgene. Dotted circle indicates the OL cell body, while brackets indicate the breadth of a single OL. (E). *erbb2* cell type-specific CRISPR quantification. Total myelin sheath length produced by individual oligodendrocytes in animals expressing Cas9 in neurons is significantly less than in the other conditions. No significant change in myelin production occurred in animals with erbb2 knockdown in RB neurons or oligodendrocytes. Error bars represent mean ± SD. A t-test was used to assess significance and p-values adjusted for multiple comparisons. Each point represents one OL from ≥ 4 animals per category; one representative experiment shown from three replicates.

## Discussion

Our results reveal a previously unappreciated function for Nrg1 type II signals and their ErbB2 receptors in myelination of axons in the CNS. A *nrg1* type II reporter is prominently expressed by RB sensory neurons in the spinal cord, and cell type-specific CRISPR experiments indicate that *nrg1* type II signals are required in these unmyelinated RB neurons to regulate the ensheathment of neighboring neurons. *In vivo* imaging indicates that myelination is reduced, but not eliminated, in two different classes of myelinated neurons in *nrg1* type II mutants. Thus, our analysis demonstrates that *nrg1* type II signals are required in unmyelinated neurons in order to regulate the extent of myelination of other neuronal classes in the spinal cord.

Our study defines a function for *nrg1* type II signals that is distinct from prior work on Nrg1 isoforms in CNS myelination (Brinkmann et al., 2008; Taveggia et al., 2008). One prior study reported that Nrg1 type III heterozygous mutant mice have thinner myelin in the brain but not the spinal cord, identifying a potential role for Nrg1 type III signals in myelination in the CNS (Taveggia et al., 2008). Nrg1 type III signals have also been implicated in increasing myelin in response to social cues (Makinodan et al., 2012), but the exact location and mode of action of Nrg1 type III has not been defined. Our analysis of the zebrafish spinal cord demonstrated a requirement for *nrg1* type II signals, which have not been examined in published mouse knockout studies of CNS myelination. In addition, our analysis revealed that *nrg1* expression in one class of neurons affects the myelination of another— an observation enabled by the use of cell type-specific knockdown approaches and in vivo myelination reporters. Consistent with prior work (Brinkmann et al., 2008; Taveggia et al., 2008), our analysis revealed normal myelination in the spinal cord of *nrg1* type III mutants and *nrg1* type I mutants, but future studies of isoform-specific mutants may investigate possible roles for these signals in other areas of the CNS.

The mode of action that we define for Nrg1 type II proteins in the spinal cord is also quite distinct from the well-studied function of Nrg1 type III proteins in the PNS. In the PNS, Nrg1 type III is essential for many steps of Schwann cell development and myelination, including proliferation, migration, radial sorting, and myelin formation (Michailov et al., 2004; Nave and Salzer, 2006; Perlin et al., 2011; Raphael et al., 2011). Whereas *Nrg1* type III mutants lack almost all myelin in the PNS, our analysis shows that myelination is only partly reduced in the spinal cord of *nrg1* type II mutants. Thus, our evidence indicates that *nrg1* type II modulates CNS myelination. Functional analyses of ErbB2 receptors reveal another important difference between the CNS and PNS: in the PNS axonal Nrg1 type

III signals activate ErbB receptors on closely associated Schwann cells, whereas in the spinal cord we show that myelination requires ErbB2 receptors not in oligodendrocytes, but in other neurons. Despite these differences, Nrg1 signals appear to exert exquisite control on myelination in both the CNS and the PNS; in both cases heterozygous mutants myelinate in proportion to Nrg1 dosage, indicating marked sensitivity to the levels of Nrg1 signals (Brinkmann et al., 2008; Michailov et al., 2004; Taveggia et al., 2008).

Based on *nrg1* type II reporter expression and our cell-specific CRISPR experiments, we propose that Nrg1 type II proteins expressed by RB neurons activate ErbB2 receptors on other neurons. Knockdown of *nrg1* type II signals in RB neurons produces a similar phenotype to loss of *erbb2* function in other neurons, suggesting that ErbB2 may be the main receptor transducing Nrg1 type II signals in this context. ErbB2 does not directly bind to Nrg1 proteins (Sliwkowski et al., 1994; Tzahar et al., 1994), so it likely that the active receptor is a heterodimer of ErbB2 with either ErbB3 or ErbB4. Although our results highlight a role for *erbb2* in neuron-to-neuron signaling in myelination, we do not discount the possibility the Nrg1 type II signals might act directly on oligodendrocytes as well, through other ErbB receptor combinations.

A key outstanding question is the mechanism by which Nrg1 type II signals are transmitted to promote spinal cord myelination. Our results indicate that dorsally located sources of Nrg1 type II signals (i.e. RB neurons) promote myelination of both dorsal and ventral axons that project from distantly located neurons in the hindbrain and spinal cord. Because the Ig-like domain of Nrg1 II binds extracellular matrix at synapses in other contexts, limiting diffusion and potentially increasing the signaling potency (Li and Loeb, 2001; Loeb et al., 1999; Pankonin et al., 2005), it seems likely that Nrg1 type II acts to promote myelination via a short-range mechanism. For example, RB, CoPA, and RS neurons all participate in the spinal cord escape response circuit, and it is possible that RB neurons express Nrg1 type II signals to regulate the myelination status of their synaptic partners. Touch activates RB sensory neurons to trigger motor neuron activation and a characteristic escape behavior (Drapeau et al., 2002; Faber et al., 1989). RB axons ascend to the hindbrain, where RS cell bodies reside (Bernhardt et al., 1990; Metcalfe et al., 1990) and are activated in response to touch on the trunk (Liu and Fetcho, 1999; O’ Malley et al., 1996). RB axons also synapse onto CoPA neurons in the dorsal spinal cord (Bernhardt et al., 1990; Gleason et al., 2003; Hale et al., 2001; Pietri et al., 2009) where they likely modulate RB sensory input (Knogler and Drapeau, 2014; Pietri et al., 2009). Thus, as the circuit forms, Nrg1 type II from RB neurons could signal RS and CoPA neurons to increase their myelination. In various other contexts, myelination is modulated to coordinate the output of circuits of disparate length: examples occur in the cerebellum, retina, and fish electromotor systems (Bennett, 1970; Stanford, 1987; Sugihara et al., 1993; Waxman, 1997). RB neurons along the spinal cord project varying distances to their RS targets in the hindbrain, so perhaps Nrg1 type II signals modulate myelination to optimize escape response circuits.

This study provides evidence that Nrg1 type II signals modulate myelination of distinct neuronal classes in the spinal cord. Our *nrg1* type II reporter and cell-specific CRISPR experiments support a model in which unmyelinated neurons express Nrg1 type II signals to regulate myelination of circuit partners, a mode of action that may coordinate function of circuits in the CNS involving both unmyelinated and myelinated neurons.

## Supporting information

Key Resources Table

## Acknowledgements

This work was supported by the National Institutes of Health [R35 NS111584 to W.S.T., 1F32NS095466 to D.E.L.]. We thank members of our labs for helpful discussion and critical comments on the manuscript and Tuky Reyes and Chenelle Hill for fish care. W.S.T. is a Kennedy-Grossman Fellow in Human Biology at Stanford University. We are grateful for the kind gift of the GFP2AtdTomato-cntn dual color myelin reporter and cell-specific CRISPR constructs from Rafael Almeida and David Lyons. We are grateful to the labs of Alvaro Sagasti and Kelly Monk for sending constructs, and to Jacob Hines and Kristin Bruk Artinger for helpful discussion.

## Author contributions

DEL: Investigation, Formal analysis, Software, Validation, Data Curation, Visualization, Writing – Original Draft

WST: Resources, Supervision, Project Administration

DEL and WST: Conceptualization, Methodology, Writing – Review & Editing, Funding acquisition

## Declaration of Interests

There authors have no competing interests to declare.

---

**Figure S1. *mbp* is expressed in the CNS of *nrg1* null and isoform-specific mutants *In situ* hybridization showing *mbp* expression by myelinating glia in the PNS (arrowheads) and CNS (arrow)**. (A) Wildtype animals show prominent expression along the lateral line nerve (arrowheads) and in the spinal cord. (B) *erbb2*^st61^ mutants, (C) *nrg1* EGF mutants, and (D) *nrg1* type III mutants all lack expression of *mbp* in the PNS, but still maintain *mbp* expression in the CNS. (E) *nrg1* type I mutants and (F) *nrg1* type II mutants maintain *mbp* expression in both PNS and CNS. (G) Oligodendrocyte numbers appear normal in the dorsal spinal cord in animals lacking EGF and *nrg1* type I and II, with a slight reduction in animals lacking *nrg1* type III. Mann-Whitney test.

**Table S1**. Cloning and genotyping primers, sgRNA sequences

## METHODS

See Key Resources Table for zebrafish lines and constructs used in this study.

## RESOURCE AVAILABILITY

### Lead contact

Further information and requests for resources and reagents should be directed to and will be fulfilled by the lead contact, William Talbot.

### Materials availability

Reagents generated in this study will be made available on request to the Lead contact. Data and code availability

DNA sequence information and microscopy images are available upon request. This paper does not report original code. Any additional information required to repeat experiments are available from the Lead contact upon request.

## EXPERIMENTAL MODEL AND SUBJECT DETAILS

### Zebrafish Lines and Maintenance

All zebrafish experiments were conducted under protocols approved by the Stanford University institutional animal care and use committee and conforming to appropriate city, state, and national regulations. Prior to experimental procedures embryos and larvae were anesthetized with 0.016% Tricaine. To inhibit pigmentation, embryos and larvae were treated with 1-phenyl 2-thiourea (PTU, 0.003%). Embryos and larvae were analyzed up to 6 dpf, before the onset of sexual differentiation.

## METHOD DETAILS

### *nrg1* allele generation and genotyping

The *nrg1* alleles st150, st151, st152, and st153 were generated by injecting sgRNAs (Table S1) and Cas9 mRNA into wildtype embryos at the one-cell stage. Injected fish were raised to adulthood and outcrossed to establish stable lines bearing variants in *nrg1*. The *nrg1* type III st153 allele has been previously described (Lysko et al., 2022). The first EGF exon common to all known isoforms was targeted to create st150; a 4 bp deletion in the EGF domain that introduces a frameshift and premature stop (P264R_fs*270) predicted to disrupt the function of all *nrg1* isoforms. Similar approaches were used to target isoform-specific exons, as illustrated in Figure 1: (st151: 29 bp deletion, A6E_fs*8), (st152: 31 bp deletion, V44C_fs*63), (st153: 7 bp deletion, P40T_fs*68). The larger deletions st151 (*nrg1* type I) and st152 (nrg1 type II) were readily genotyped using primers (Table S1) that generate a shorter product from the mutant allele that is distinguishable from the wildtype allele on an agarose gel. st150 disrupts a BsaJI restriction enzyme site; PCR primers oAMS_618/619 amplify a 430 bp fragment from the wildtype allele that BsaJ1 cleaves into three fragments (208, 158 and 64 bp), while the mutant product is cleaved into two fragments (218, 208 bp). st153 disrupts a BslI site; primers CRD_L2/R2 results amplify a 488 bp fragment from the mutant allele that is cleaved by BslI into three fragments (304, 99, 77 bp), whereas the wildtype product is cleaved into four fragments (235, 99, 77 and 77 bp). We also identified additional frameshift alleles caused by the same guide RNAs, causing the same phenotypes (data not shown). Genotyping primers are described in Table S1.

### *In situ* hybridization

*In situ* hybridization for *mbp* mRNA in whole-mount larvae was performed using standard methods as described (Lyons et al., 2005; Thisse and Thisse, 2014).

### Oligodendrocyte sheath analysis

The stable transgenic line claudink:Gal4-UAS:GFP-CAAX (Münzel et al., 2012) was crossed to *nrg1* mutant lines and used to visualize the myelin sheaths of individual oligodendrocytes at 3.5 dpf. At this stage, a number of oligodendrocytes have migrated to the dorsal region of the spinal cord and many have completed the process of myelin sheath initiation and maturation (Czopka et al., 2013). In this dorsal region, all the sheaths generated by an oligodendrocyte can be visualized and analyzed discretely from neighboring oligodendrocytes. For sheath analysis at 5 dpf, we injected mbp:GFP-CAAX (Almeida et al., 2011) to scatter-label individual oligodendrocytes in transiently transgenic animals. We imaged live larvae anaesthetized and embedded in agarose using a Zeiss LSM700 confocal microscope. Sheaths belonging to an individual oligodendrocyte were quantitated in ImageJ by drawing a segmented line along the length of each sheath; a script in Rstudio collated sheath parameters per cell.

### Antifreeze protein 3’ UTR construct cloning

p3E-afp_UTR was created and used as a 3’ Entry clone for LR-generated constructs in this study. Sequence encoding the ocean pout antifreeze (afp) protein 3’ UTR (Horstick et al., 2015) was ordered as a gBlock (IDT) and cloned into pDonP2R-P3 using Gibson cloning (NEB).

### Myelin reporter analysis

The GFP-cntn1a and GFP-2A-tdTcntn1a myelin reporters have been previously used to visualize myelin sheaths on individual neurons (Almeida et al., 2021; Koudelka et al., 2016). We used a multiple-vector strategy in transient transgenic animals in order to visualize neuronal type and myelination status in our stable *nrg1* mutant lines. We created pTol2_6XUAS:GFP-cntn1a (pDEL73) using LR cloning, and received the kind gift of pTol2_10XUAS:GFP-2A-tdTcntn1a (Almeida et al., 2021) to express the contactin myelin reporter. To drive pan-neuronal expression of these vectors we created pTol2_NBT:Kal4 (pDEL72) and pTol2_HuC:Kal4 (pDEL102) using LR cloning. To randomly label neurons we injected 0.06 ng each of a Kal4 driver vector and a myelin reporter vector into embryos at the one-cell stage. Neuronal type and myelin sheaths were imaged at 5.5-6 dpf using a Zeiss LSM700, and imaged larvae genotyped. Genotype, neuronal type, and myelination status were determined, and data were collated and analyzed using scripts in RStudio.

### *nrg1* type II-reporter transgene

Using zebrafish genomic DNA prepared using DNeasy Blood & Tissue Kits (Qiagen) as a template, we amplified two overlapping fragments covering approximately 3kb of DNA upstream from the *nrg1* type II translational initiation codon using KAPA polymerase (Roche), and the following primer pairs: (T2_3055_G_F1; 1677_R) and (1399_F; T2_G_R8). Fragments were joined using overlap-extension PCR and inserted into pDonrP4-P1R using Gibson cloning (NEBuilder HiFi) to create a p5E Gateway vector (pDEL61). Using LR cloning we created the Tol2 vector pDEL63, in which sequences from upstream of *nrg1 type II* drive expression of membrane bound GFP, with a cmlc2:GFP marker of transgenesis. Tol2 mRNA and pDEL63 were injected into embryos at the one cell stage, and animals with GFP signal in the heart were raised to adulthood. Stably expressing lines were established from the F0 generation and screened for GFP expression outside of the heart.

### Rohon-Beard-specific regulatory element LSEC

A short section of the lateral stripe element (LSE) upstream of the coding region of the *neurogenin1* gene previously shown (Blader et al., 2003) to drive expression in developing Rohon-Beard neurons was amplified with the following primer pair: (ng1_LSE_C_F1;ng1_LSE_C_F1). This fragment was inserted into pDonrP4-P1R as above to create p5E_ngn1_LSE_C (pDEL98; hereafter referred to as LSEC, describing a minimal “critical” region expressed in RBs). pME_GFP-CAAX was generated using Infusion cloning (Takara) and combined with pDEL98 to create the expression vector pTol2_LSEC:GFP-CAAX, used to assess the strength and specificity of LSEC expression.

### Cell-type specific CRISPR

We used previously described Gateway-cloning compatible Tol2 constructs that express Cas9 using a tissue-specific promoter and gene-specific gRNAs using the U6 promoter; cmlc2:GFP was included as a marker of transgenesis (Ablain et al., 2015; Marshall-Phelps et al., 2020). sgRNAs are described in Table S1; *nrg1* sgRNAs are the same as those used to generate the stable lines, with the exception of *nrg1* type III. A new gRNA (coDEL_161) for nrg1 type III was designed to avoid a SNP in the previously targeted type III-specific sequence that might lower cutting efficiency. Four *erbb2* sgRNAs were designed and evaluated for efficiency by injecting sgRNA and Cas9 protein (UC Berkeley) into wildtype embryos at the one-cell stage and assessing *mbp* expression along the peripheral lateral line nerve by *in situ* hybridization at 4.5 dpf; the most efficient sgRNA (coDEL_171) was used in subsequent experiments. Complementary oligos encoding the sgRNAs were annealed and cloned into the BseRI site of the cell-specific CRISPR destination vector using ligase (NEB). We used the LSEC element to drive expression of Cas9 in Rohon-Beard neurons (Blader et al., 2003), the NBT regulatory element to drive expression pan-neuronally (Bonner et al., 2012), and the *claudink* regulatory element to drive expression in oligodendrocytes (Münzel et al., 2012). LSEC:*nrg1* targeted Tol2 expression vectors (pDEL125-129) and *erbb2*-targeted expression vectors (pDEL133-135; pDEL142-143) were created using LR cloning as above. To analyze the effect on myelin sheaths, we injected cell-specific CRISPR constructs and Tol2 mRNA early during the one-cell stage into cldnk:GFP stable transgenic embryos, selected fish with efficient construct expression (GFP in heart) at 3.5 dpf, and imaged myelin sheaths as above.

## QUANTIFICATION AND STATISTICAL ANALYSIS

Graphs and statistical analysis were done in GraphPad Prism. Data were first tested for normality using D’ Agostino & Pearson, Anderson-Darling, and Shapiro-Wilk normality tests. If normal, comparison of means was done with ANOVA and t test for pairwise comparisons, adjusting the p value in the case of multiple comparisons. Non-normal groups were compared using the Mann-Whitney test. For comparison of proportions of myelinated axons, Fisher’ s exact test was used. Exact p-values are reported. Error bars indicate mean ± standard deviation.

## Notes

### Competing Interest Statement

The authors have declared no competing interest.

