## Supplementary material for "Unmyelinated neurons use Neuregulin signals to promote myelination of neighboring neurons in the CNS": Figure S1

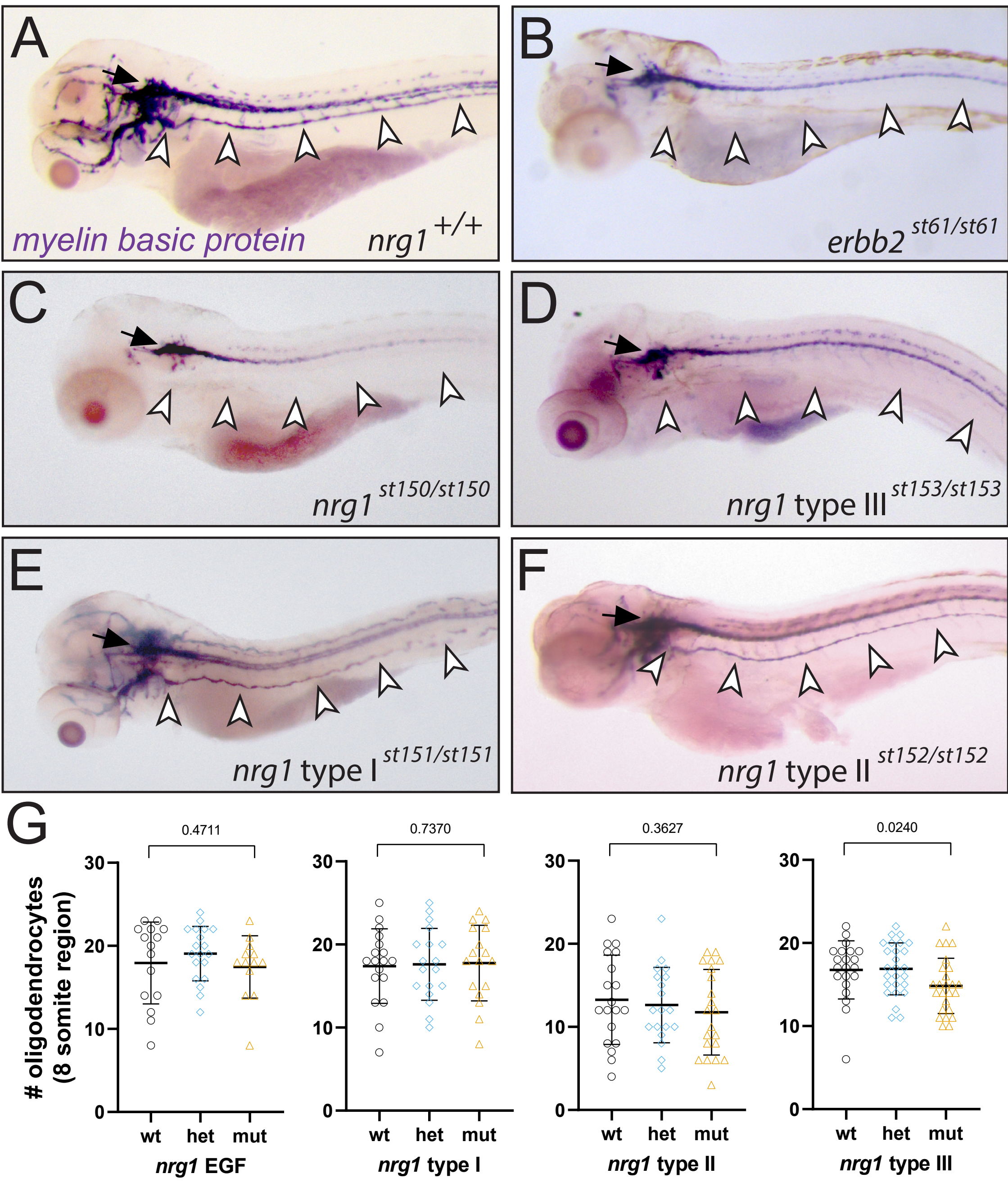
A
myelin basic protein
nrg1 +/+
B
erbb2 st61/st61
C
nrg1 st150/st150
D
nrg1 type III st153/st153
E
nrg1 type I st151/st151
F
nrg1 type II st152/st152
G
### oligodendrocytes (8 somite region)
0.4711
0.7370
0.3627
0.0240
30
20
10
0
wt
het
mut
nrg1 EGF
nrg1 type I
nrg1 type II
nrg1 type III

**Figure S1. *mbp* is expressed in the CNS of *nrg1* null and isoform-specific mutants** *In situ* hybridization showing *mbp* expression by myelinating glia in the PNS (arrowheads) and CNS (arrow). (A) Wildtype animals show prominent expression along the lateral line nerve (arrowheads) and in the spinal cord. (B) *erbb2*$^{st61}$ mutants, (C) *nrg1* EGF mutants, and (D) *nrg1* type III mutants all lack expression of *mbp* in the PNS, but still maintain *mbp* expression in the CNS. (E) *nrg1* type I mutants and (F) *nrg1* type II mutants maintain *mbp* expression in both PNS and CNS. (G) Oligodendrocyte numbers appear normal in the dorsal spinal cord in animals lacking EGF and *nrg1* type I and II, with a slight reduction in animals lacking *nrg1* type III. Mann-Whitney test.
