## Supplementary material for "Unmyelinated neurons use Neuregulin signals to promote myelination of neighboring neurons in the CNS": Table S1

**Table S1.** Cloning and genotyping primers, sgRNA sequences

| Primer Name | Use | Sequence |
|---|---|---|
| T2_3055_G_F1 | nrg1 type II -3kb amp | CCCATCGATTCGAATTCCCACTGAGCCACCATGTAGC |
| 1677_R | nrg1 type II -3kb amp | GCTGAACTCTGCATGAAAGGG |
| 1399_F | nrg1 type II -3kb amp | CCCTTTCATGCAGAGTTCAGC |
| T2_G_R8 | nrg1 type II -3kb amp | ACCGGTTCTAGAGGCTCCTCCGCGCTCGG |
| P4_P1R_F1 | pDONR_P4-P1R_amp | GAGCCTCTAGAACCGGTGG |
| P4_P1R_R1 | pDONR_P4-P1R_amp | GAATTCGAATCGATGGGATCC |
| ng1_LSE_C_F1 | neurogenin1 LSEC amp | CCCATCGATTCGAATTCACTACTTTCCTTCCCCCTCTCC |
| ng1_LSE_C_R1 | neurogenin1 LSEC amp | ACCGGTTCTAGAGGCTCGGGAAAGAAGTCTGAGCTTGACTC |
| **sgRNA name** | **Use** | **Sequence** |
| coAMS 02 | target nrg1 EGF | GGATGTCTTGGCCGAGGGAGTGG |
| coDEL_02 | target nrg1 type I | GCAGGCAAAGAAGGGAAAAGTGG |
| coDEL_42 | target nrg1 type II | GGAGCTCGCCCGCCGCTCGGGGG |
| coDEL_60 | target nrg1 type III | GGAGAGAACGTGCCGGGGGACGG |
| coDEL_161 | target nrg1 type III | GGGACGGTGACAGCTCCGAGCGG |
| coDEL_171 | target erbb2 | CCAACCTTCTCTCCCTGTGGCGG |
| **Genotyping primers** | **Use** | **Sequence** |
| oAMS_618 | st150 genotyping | GGTAGATGAAATGCTGTTCACTC |
| oAMS_619 | st150 genotyping | GCGCATAGTTTGCATGTC |
| oDEL_2 | st151 genotyping | GGAGTCCACATTCACACTGCG |
| oDEL_1 | st151 genotyping | CCTTGCGGAGAGTTGTCAGG |
| T2_F3 | st152 genotyping | GGTCTCGGTTCTTTACGGAGC |
| T2_R5 | st152 genotyping | GCTCTTCCTCCTGGAGCTTCC |
| CRD L2 | st153 genotyping | GGATCCTGTCCTATACGCTTT |
| CRD R2 | st153 genotyping | CGCTGCTGATCTGATAATGG |
