## Supplementary material for "Unmyelinated neurons use Neuregulin signals to promote myelination of neighboring neurons in the CNS": Key Resources Table

| REAGENT or RESOURCE | SOURCE | IDENTIFIER |
|---|---|---|
| **Experimental models: Organisms/strains** | | |
| nrg1 EGF | This paper | st150 |
| nrg1 type I | This paper | st151 |
| nrg1 type II | This paper | st152 |
| nrg1 type III | (Lysko et al. 2022) | st153 |
| Tg([-3kb]nrg1 typeII:GFP-CAAX) | This paper | st162 |
| Tg(claudink:GAL4-UAS:GFP-CAAX) | (Münzel et al. 2012) | |
| Tg(NBT:dsRed) | (Peri and Nüsslein-Volhard 2008) | zf148Tg |
| **Oligonucleotides** | | |
| For all oligonucleotides used, see Table S1 | This paper | NA |
| **Recombinant DNA** | | |
| pTol2_mbp:GFP-CAAX | (Almeida et al. 2011) | |
| p5E-NBT | (Peri and Nüsslein-Volhard 2008) | |
| p5E-HuC | (Shiau et al. 2013) | |
| pME-KalTA4 | (Almeida and Lyons 2015) | |
| p3E-afp_UTR | This paper | pAMS448 |
| Tol2kit | (Kwan et al. 2007) | |
| pTol2_NBT:Kal4 | This paper | pDEL72 |
| pTol2_HuC:Kal4 | This paper | pDEL102 |
| p5E-6XUAS | (Almeida and Lyons 2015) | |
| pCR8/GTW-GFP_cntn1a | (Koudelka et al. 2016) | |
| pTol2_6XUAS:GFP-cntn | This paper | pDEL73 |
| pTol2_6XUAS:tdT-cntn | This paper | pDEL74 |
| pTol2_10UAS-EGFP-2A-zftdTomato-cntn1a_crymCherry | (Almeida et al. 2021) | |
| p5E_(-3kb)nrg1 type II | This paper | pDEL61 |
| pTol2_(-3kb)nrg1_typeII:GFP-CAAX | This paper | pDEL63 |
| p5E_claudinK | (Münzel et al. 2012) | |
| p5E_ngn1_LSE_C | This paper | pDEL98 |
| pME_GFP-CAAX | This paper | pELB01 |
| pTol2_LSEC:GFP-CAAX | This paper | pDEL116 |
| pME_memtagRFPt T2A cas9 | (Marshall-Phelps et al. 2020) | |
| pME_cas9_2A_GFP | Addgene | #63155 |
| pDest_U6_A2_CG2 | Addgene | #63156 |
| pDest_U6_A2_CG2_typeEGF_gRNA | This paper | pDEL110 |
| pDest_U6_A2_CG2_T1_gRNA | This paper | pDEL111 |
| pDest_U6_A2_CG2_T2_gRNA | This paper | pDEL112 |
| pDest_U6_A2_CG2_T3_gRNA | This paper | pDEL113 |
| pTol2_ngn1:redCas9; empty_sgRNA (LSEC) | This paper | pDEL127 |
| pTol2_ngn1:Cas9green; EGF_sgRNA (LSEC) | This paper | pDEL128 |
| pTol2_ngn1:Cas9green; T1_sgRNA (LSEC) | This paper | pDEL126 |
| pTol2_ngn1:Cas9green; T2_sgRNA (LSEC) | This paper | pDEL129 |
| pTol2_ngn1:Cas9green; T3_sgRNA (LSEC) | This paper | pDEL125 |
| pDest_U6_A2_CG2_erbb2_gRNA171 | This paper | pDEL133 |
| pTol2_claudink:Cas9green; empty_gRNA | This paper | pDEL143 |
| pTol2_NBT:Cas9green; erbb2_gRNA171 | This paper | pDEL134 |
| pTol2_LSEC:Cas9green; erbb2_gRNA171 | This paper | pDEL135 |
| pTol2_claudink:Cas9green; erbb2_gRNA171 | This paper | pDEL142 |
| **Software and algorithms** | | |
| Fiji | (Schindelin et al. 2012) | RRID: SCR_002285 |
| GraphPad Prism | GraphPad Software | RRID:SCR_002798 |
| Adobe Illustrator | Adobe | RRID: SCR_010279 |
| Rstudio | RStudio | RRID:SCR_000432 |
| R | R Project for Statistical Computing | RRID:SCR_001905 |
